# Ultrafast Frame-Free Imaging of Neural Activity with Event Cameras

**DOI:** 10.64898/2026.01.08.698496

**Authors:** Angelo Forli, Krishna Chaitanya Kasuba, Katharine Henn, Deborah Lee, Miriam Hernández-Morales, Chunlei Liu, Evan W. Miller, Michael M. Yartsev

**Author notes:** These authors contributed equally (Angelo Forli, Krishna Chaitanya Kasuba).

## Abstract

Frame-based fluorescence imaging has long defined how neural activity is optically measured. This approach requires acquiring all pixels within an image, regardless of whether they carry meaningful neural dynamics, thereby intrinsically coupling spatial and temporal resolution while increasing data output. Here, we introduce an entirely different, frame-free approach that leverages the sparse nature of neural activity using event-based cameras, which asynchronously report fluorescence changes as spatiotemporal events. Compared with a frame-based camera, our method preserves signal fidelity while eliminating the fixed trade-off between spatial resolution, temporal resolution and data rate, thereby reducing data output by orders of magnitude. Applied to hippocampal preparations we demonstrate that the frame-free approach can resolve both single action potentials and fast network dynamics over large fields of view at kilohertz rates, enabling scalable, ultrafast optical recordings.

## Main

Traditional approaches for imaging functional neural activity, such as calcium and voltage imaging, have transformed our understanding of brain function by enabling single-cell recordings across large neural populations in behaving animals^1^. However, these approaches largely rely on frame-based cameras (FCs), which synchronously acquire all pixels irrespective of whether they carry informative fluorescence dynamics. For neural signals that are often sparse in space and time^2^ this acquisition paradigm can be inefficient, leading to rapidly escalating data rates, as frame rate and field of view increase. In practice, limitations in bandwidth, power consumption, heat dissipation, and hardware footprint intrinsically couple spatiotemporal resolution and data rate^1,3^, often necessitating sophisticated optical or computational strategies^4–6^ to mitigate these trade-offs. This problem is especially acute for voltage imaging, limiting the development of microscopes for full-frame acquisition at kilohertz rates while maintaining high spatial resolution over a large field of view^1^ (FOV), as well as of miniaturized, untethered microscopes for high-speed imaging in freely moving animals^7^.

Here, we propose a new approach for addressing these challenges using a new type of camera that has yet to be utilized for imaging neural activity: the event-based camera (EC, **Fig. 1a**). ECs are based on neuromorphic sensors and radically differ from traditional FCs in their operation. Rather than capturing frames at a fixed rate, they asynchronously detect changes in brightness at each pixel, generating a continuous stream of events that encode the time, location, and polarity (positive or negative) of these changes^8^. Compared to conventional cameras, ECs offer several advantages: exceptional temporal resolution and low latency (both at the µs scale), extensive dynamic range, minimal power consumption and reduced data output. These novel sensors have shown success in various applications^8,9^, including mobile robotics, ultrafast particle tracking and super-resolution microscopy^10,11^. Yet, despite their vast potential for fast, large-scale and scalable neural activity imaging, the use of ECs for directly capturing biologically meaningful neural signals, such as calcium or voltage in neurons, remains virtually unexplored^12,13^.

**Figure 1.**
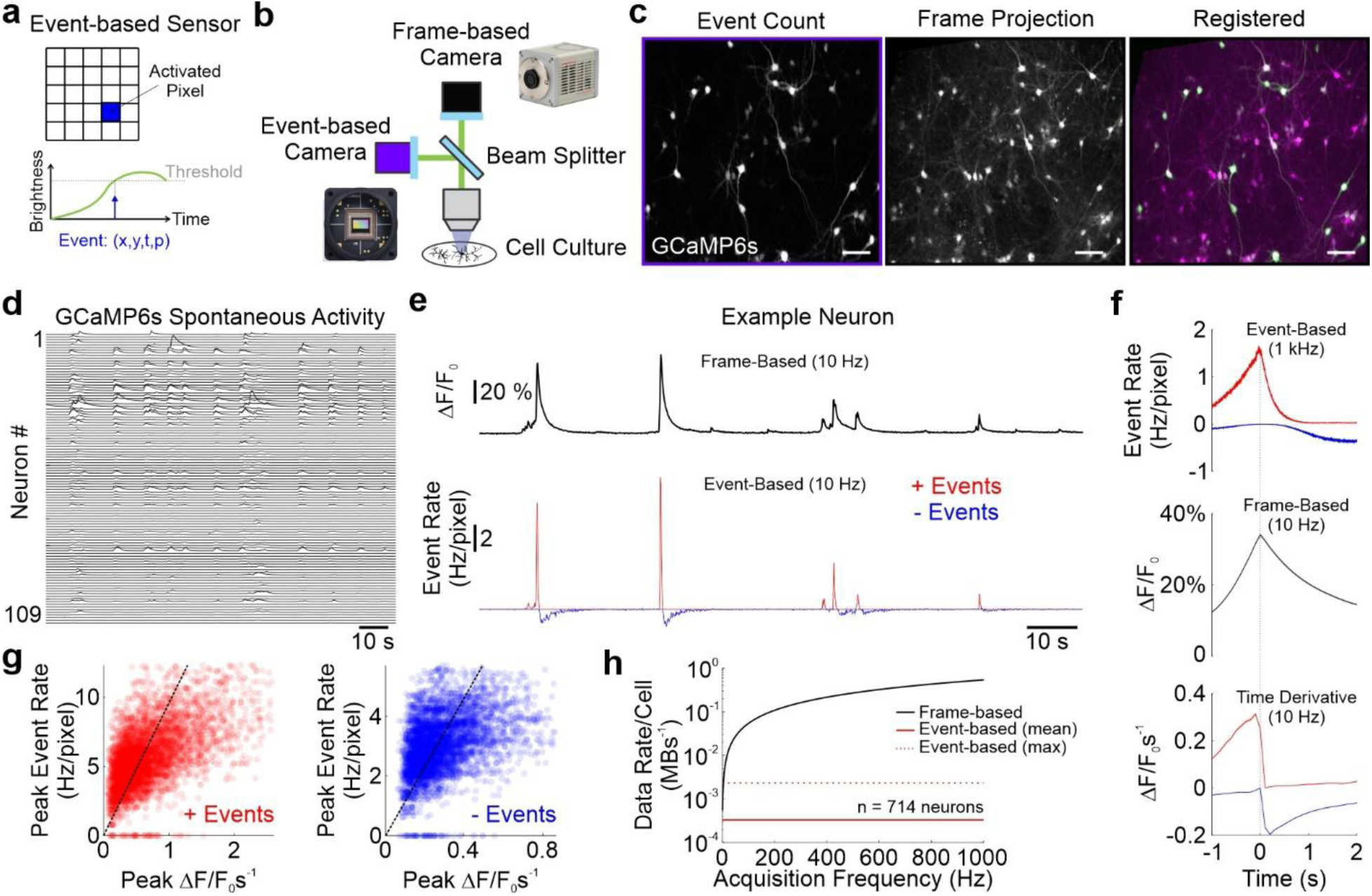
Neural activity imaging with an event-based camera. **a,** Representation of operation of an event-based sensor. Pixels are activated (blue) asynchronously when the brightness crosses an adjustable threshold (Methods), generating an event (*x* and *y*: pixel coordinates; *t*: time; *p*: polarity, ± 1 of the event). **b**, Microscope schematics and pictures of the cameras used in this study. **c**, *Left*: grayscale image representing the event count distribution across EC pixels, obtained from a representative recording. *Center*: average pixel intensity across all frames obtained from the same recording with a FC. *Right*: overlayed images after registration. Scale bars: 50 µm. **d**, Fluorescence time series (ΔF/F_0_, Methods) from simultaneously imaged ROIs acquired with a FC during spontaneous activity *in vitro*. **e**, FC ΔF/F_0_ (top) vs. EC event rate (bottom) traces for the same example ROI. Note that since FC acquisition rate was set at 10 Hz, event rate was calculated on 100 ms time bins for signal visualization at the same sampling frequency. **f**, *Top*: average event rate at 1 kHz sampling for positive (red) or negative (blue) events recorded during identified GCaMP fluorescence transients (n = 714 cells, N=6 coverslips prepared from two neuronal cultures, Methods). *Middle*: average ΔF/F_0_ during the same GCaMP transients. Bottom: Positive (red) or negative (blue) time derivative of the ΔF/F_0_ during the same GCaMP transients (Methods). Note how the event rate matches the ΔF/F_0_ time derivative. **g**, Scatter plots showing for each detected GCaMP fluorescence transient the peak positive (red) or negative (blue) event rate vs. the peak ΔF/F_0_ time derivative. Dashed lines represent the best fit. **h**, Experimentally recorded data rate per cell as a function of the acquisition frequency for the FC (black trace) and for the EC (red traces, solid: average across neurons; dotted: value for the most active neuron).

To test the feasibility of using ECs for imaging neural activity we built an epifluorescence microscope (**Fig. 1b**) equipped with both a state-of-the-art frame-based camera (Hamamatsu Orca Flash 4.0 V3) and a commercially available event camera (Sony-Prophesee IMX636) to enable side-by-side comparison. A beamsplitter in the detection path separates light from the sample such that both cameras acquire a fraction of the same signal, allowing direct comparison of their performances. We first benchmarked the microscope by imaging spontaneous neural activity *in vitro* from hippocampal neuron cultures expressing the calcium indicator GCaMP6s^14^ (N = 6 coverslips from two cultures). The FOV of the FC was binned 4x (512 x 512 pixels) to remain within reasonable data rates, whereas full resolution (1280 x 720 pixels) was preserved for the EC. The EC output is inherently frameless, consisting of a list of events reporting the timestamp, location and polarity of the corresponding brightness change. From this data stream, a static image of the sample could be obtained by quantifying the event count per pixel (**Fig. 1c**, left), representing the regions with fluorescence dynamics in the FOV, *i.e.*, neurons exhibiting calcium transients. We first registered this image with the average projection of the FC time series acquired at 10 Hz (**Fig. 1c**, center and right, Methods), such that each region of the EC FOV could be mapped to the corresponding region of the FC FOV. Next, we extracted fluorescence activity traces (ΔF/F_0_) from automatically segmented ROIs corresponding to the somata of active neurons (**Fig. 1d**). For each corresponding ROIs in the EC data, positive and negative *event rate* traces were computed separately as the sum of positive (or negative) events across all pixels within the ROI in consecutive time bins, divided by the bin interval (example ROI in **Fig. 1e**). Comparison of ΔF/F_0_ vs. the event rate for the same ROI confirmed the ability of the EC to reliably capture neural dynamics at high signal-to-noise ratio. To quantify the relationship between events and GCaMP fluorescence signals corresponding to neural activity, we detected all GCaMP fluorescence transients (Methods) and compared the average time course of the event rate vs. either the ΔF/F_0_ (**Fig. 1f**, top vs middle) or the signed time derivative of ΔF/F_0_ (**Fig. 1f**, top vs bottom). As expected from the EC sensitivity to brightness changes, the positive event rate closely matched the ΔF/F_0_ positive derivative, and their peak amplitudes were linearly related (**Fig. 1f-g**, red, Pearson *C* = 0.57, n = 5159 fluorescence transients). A similar trend was observed for the negative event rate, albeit with substantial delay (**Fig. 1f-g**, blue, Pearson *C* = 0.35, n = 5159 fluorescence transients).

A key advantage of event-based acquisition is its reduced data output, which alleviates bandwidth constraints while also relaxing downstream requirements on data storage, transmission, and power consumption. To quantify the empirical reduction in data rate, we examined the average rate of events per cell recorded during spontaneous GCaMP activity imaging (n = 714 neurons, N=6 coverslips from two cultures) and compared the corresponding data rate of the EC (Methods) with that of FC imaging the same neurons at increasing frame rates (**Fig. 1h**), assuming the same number of pixels per cell. Under our conditions (data rate would depend on activity levels), we observed on average more than two orders of magnitude data reduction at > 100 Hz acquisition frequency, with a data rate *per* active neuron well below 10^-3^ MBs^-1^ for the EC, and 0.1-1 MBs^-1^ for the FC.

Another advantage of the EC is that the time interval for binning the event rate is completely arbitrary (up to a few kHz) and does not need to be defined during the acquisition, effectively allowing a full decoupling of the spatial and temporal resolution. We leveraged this property by examining spontaneous and fast neural dynamics across large FOVs at full frame resolution (1280 x 720 pixels), by investigating the existence of repeatable neural activation sequences *in vitro* (Methods). Importantly, we did so by relying uniquely on the EC data, both for cell segmentation from high resolution even count distributions (**Fig. 2a**) and for neural activity quantification at kHz temporal resolution (**Fig. 2b**). Analysis of spontaneous activity epochs from segmented neurons revealed the existence of repeatable activation sequences in several FOVs (n=6), where large numbers of neurons were simultaneously active within a short time interval with a consistent order (*p* < 0.05, Methods), akin to neuronal sequences observed *in vivo*^15,16^ and *in vitro*^17,18^. Here, by imaging neural dynamics at megapixel resolution and kilohertz rates, we demonstrate a regime that is effectively inaccessible to conventional FCs, where spatial and temporal resolution are intrinsically constrained by data rate.

**Figure 2.**
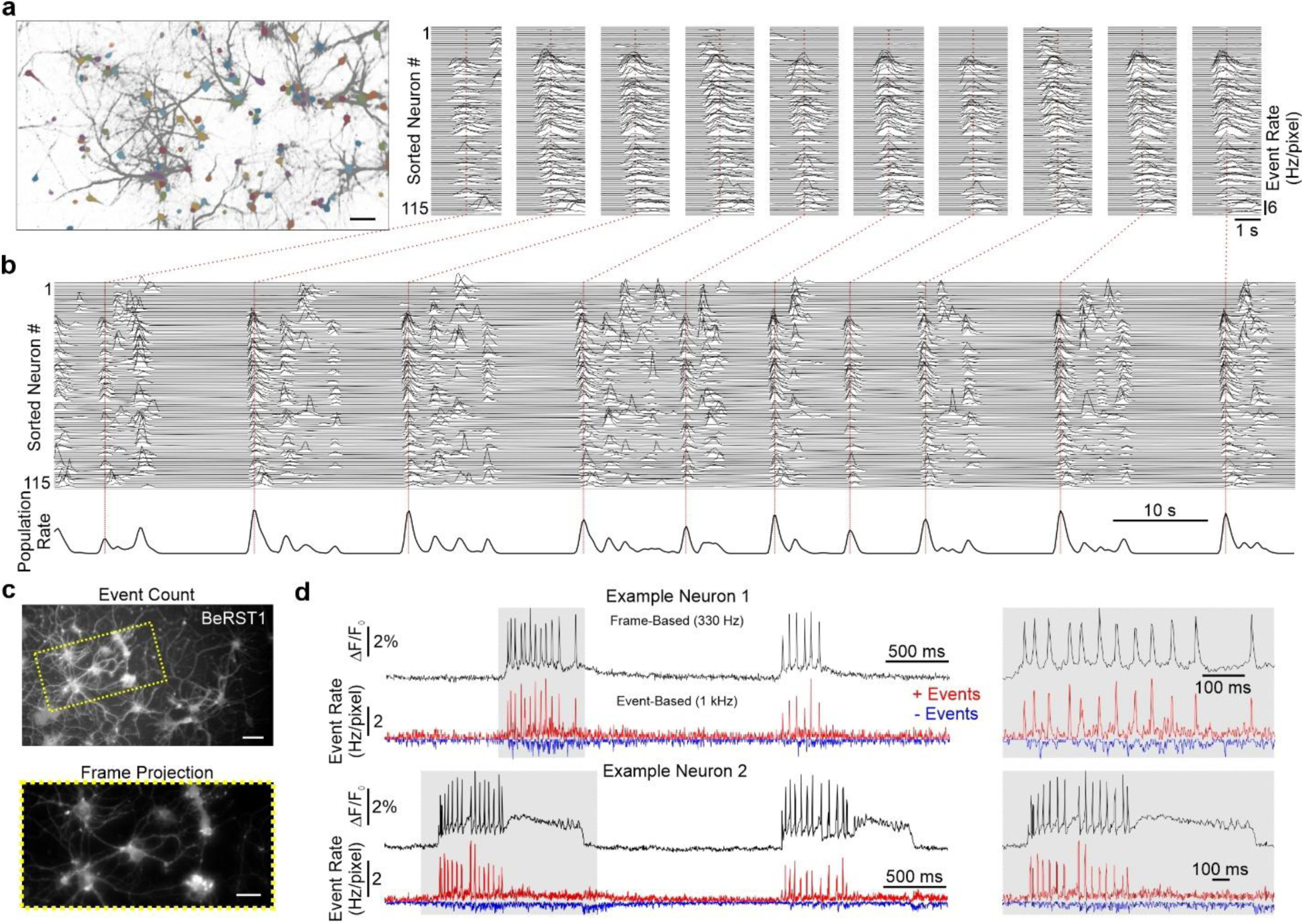
Ultrafast calcium imaging at megapixel resolution and voltage imaging of single action potentials with event-cameras. **a,** Representative image of detected events (gray points) during spontaneous GCaMP imaging *in vitro* at full resolution (1280 x 720 pixels). Different colored masks for automatically segmented ROIs (Methods) are overlaid on top. Scale bar: 50 µm. **b**, Event rate time series (1 kHz sampling) for simultaneously recorded ROIs during spontaneous activity *in vitro*. Bottom trace represents the average event rate from all the ROIs. Vertical red lines indicate detected population bursts (Methods); ROIs are sorted based on the time of activation during the burst (Methods). *Inset*: Magnified view of the event rate across imaged ROIs for each detected population burst. Note the consistent order of activation of ROIs across consecutive burst. **c**, *Top*: grayscale image representing the total event count distribution across EC pixels, obtained from a hippocampal culture loaded with BeRST1. *Bottom*: average frame projection from the FC, while imaging from the same sample. FOV indicated by the yellow rectangle. Scale bars: 20 µm **d**, *Left*: ΔF/F_0_ (black) and smoothed event rate (red and blue) traces from BeRST1 fluorescence for the same example neurons during spontaneous action potential firing *in vitro*. *Right*: magnified views of selected epochs (gray rectangles) showing the correspondence between action potential fluorescence signals imaged with the FC and the EC.

Finally, as a test case for high-speed neural activity monitoring with ECs, we turned to voltage imaging. When looking at action potentials (AP), for example, the information on whether a given neuron has fired an AP can be extracted from a relatively small fraction of pixels compared to those actively being read out at any given time. ECs are well-suited for capturing such signals, as they only capture and transmit brightness changes. However, the relatively small amplitude and short-lived nature of fluorescence voltage signals (as compared to calcium events) make them difficult to detect. To investigate the feasibility of detecting voltage signals with the EC, we loaded hippocampal cultures with the red fluorescent voltage indicator BeRST1^19^ (**Fig. 2c**) and imaged spontaneous activity *in vitro* using the same microscope equipped with both the EC and FC. Due to bandwidth limitations of the traditional FC, voltage imaging was limited to regions of 256 x 128 pixels acquired at 330 Hz, while the EC operated at full resolution. This again highlights a practical regime in which ECs enable simultaneous high temporal resolution and large field of view, a balance that is difficult to achieve with conventional frame-based cameras. Brief brightness changes, putatively associated with spontaneous single action potential firing, could be reliably detected in the cell fluorescence from the FC (**Fig. 2d**, black traces); correspondingly, the same ROIs (Methods) in the EC were associated with transient event rate changes (**Fig. 2d**, red and blue traces at 1 kHz), confirming the ability of the camera to capture small-amplitude and short-lived (ms) fluorescence transients, corresponding to single action potentials, at full spatial resolution.

In summary, we present an ultrafast, frame-free approach to neural activity imaging using ECs. The sparse and asynchronous output of ECs decouples spatial and temporal resolution, yielding orders-of-magnitude reductions in data rate while enabling kilohertz-scale imaging across large fields of view and the detection of single action potentials at full-frame resolution. Although here we focus on the direct acquisition of fast fluorescence changes, any time-varying signal can, in principle, be reconstructed by temporal integration, provided that absolute-intensity references are sampled sufficiently often to limit drift. Emerging hybrid sensors that combine event- and frame-based modalities on the same chip provide a natural pathway toward faithful reconstruction of neural signals spanning multiple temporal scales at high spatial resolution. More broadly, our results position event-based sensing as a transformative architectural foundation for neural imaging, alleviating fundamental constraints on spatiotemporal resolution, data bandwidth and power consumption, and opening new opportunities for wireless voltage imaging and the study of neural activity in freely behaving, and potentially even flying, animals.

## Methods

### Microscope

The microscope used for all experiments was a custom-made setup based on the Cerna Modular Microscopy Platform (Thorlabs). Briefly, excitation light was provided by either a blue (Thorlabs M455L4, 455 nm for GCaMP experiments) or a red (Thorlabs SOLIS-660D, 660 nm for BeRST1 experiments) LED connected to an epi-illumination path (Thorlabs WFA2001 for GCaMP or Thorlabs CSE2100 for BeRST1). Green or red fluorescence filter sets were used to separate excitation and emission light and the sample was imaged through a 10x objective (Nikon CFI Fluor 10x, 0.3 NA for GCaMP imaging) or a 20x objective (Olympus XLUMPlanFl, 20x, 0.95 NA, for BeRST1 imaging). A beamsplitter (50/50 reflected/transmitted ratio, Thorlabs BSW10R for GCaMP imaging or 70/30, Thorlabs BST10R for BeRST1 imaging) was utilized to separate fluorescent light from the sample into two paths: one directed to the EC (reflected component) and the other directed to the FC (transmitted component). The cameras (Hamamatsu Orca Flash 4.0 V3, FC; Sony-Prophesee IMX636, EC) were mounted on 1x and 0.5x camera tubes, respectively, mounted on a two camera mount (Thorlabs 2CM1), and their position finely adjusted for imaging the same focal plane. The sample was mounted on a motorized movable platform (xy) and the entire microscope mounted on an optical table. Frame start times were timestamped in the EC by connecting the frame-output port of the EC to the synch-in port of the EC, and used for synchronization during post-processing. Data from both cameras were acquired by a workstation, through USB 3.0 for the EC and a camera link (Full Configuration Deca Mode, 10 Taps) for the FC. EC data were acquired in Metavision Studio software (Prophesee) and FC data were acquired in HC Image (Hamamatsu). The EC provides a set of adjustable settings (*biases*), allowing the sensor performance to be tuned for different application requirements. Biases include on/off event thresholds, low/high pass temporal filters and a refractory period between consecutive events from the same pixel. Most recordings were obtained with default biases. A home-built 12 V Peltier device with a custom 3D printed dish holder was used to maintain the temperature of the cell culture dishes at 37 °C during imaging.

### Neuronal Cultures

Hippocampi were isolated from embryonic day 18 Sprague Dawley rat embryos (Charles River Laboratories) and transferred into ice-cold, sterile Hank’s balanced salt solution lacking Ca²⁺ and Mg²⁺. Unless noted otherwise, reagents used for dissection and culture were obtained from Invitrogen. The tissue was enzymatically dissociated by incubation in 2.5% trypsin for 15 min at 37 °C, followed by gentle trituration with fire-polished Pasteur pipettes. Cells were resuspended in minimum essential medium (MEM) supplemented with 5% fetal bovine serum (Thermo Scientific), 2% B-27, 2% (1 M) D-glucose (Fisher Scientific), and 1% GlutaMAX, and then plated onto 12-mm coverslips (Electron Microscopy Sciences; prepared as described above) at 27,000 cells per coverslip. Cultures were maintained at 37 °C in a humidified 5% CO₂ incubator. At 1 day in vitro (DIV), 50% of the MEM-based medium was exchanged for Neurobasal medium containing 2% B-27 and 1% GlutaMAX. Imaging experiments were conducted on neurons at 10–20 DIV unless otherwise indicated.

### GCaMP6 expression and recordings

Genetically encoded calcium indicators were delivered to cultured neurons via adeno-associated virus. AAV1.Syn.GCaMP6s.WPRE.SV40 (1.8 × 10¹³ vg mL⁻¹, purchased from the University of Pennsylvania Viral Vector Core) was applied at a multiplicity of infection (MOI) of 5.0 × 10⁴ to drive expression of GCaMP6S, respectively. Neurons were transduced at 2–6 days in vitro (DIV), fluorescence expression was typically detectable by 5–9 DIV, and.

### BeRST1 loading and recordings

Hippocampal neurons were loaded with BeRST1 by diluting a 1 mM DMSO stock to a final concentration of 1μM in HBSS (Gibco). Cultures were incubated in the BeRST1–HBSS solution for 20 min at 37 °C in a humidified incubator, then washed with fresh HBSS 3x times and transferred to fresh HBSS for imaging. Spontaneous activity was recorded in these neurons maintained at 37 °C.

### Data Analysis

#### Processing of EC data

Raw data from the EC consisted of a table of consecutive events reporting the time of the event (at µs resolution), the x and y pixel location (on a 1280 x 720 grid) and the polarity (± 1). To align FC and EC data in time, all event times were referenced to the timestamp of the first frame of the FC. Event count distributions across EC pixels were generated by counting the total number of events – regardless of polarity – at a given pixel, and mapping the event number to a grayscale image between 0 (black) and the maximum number of events (white) for that recording. Event rates were calculated for each ROI as follows: first, pixels belonging to a given ROI were identified (see below) and only events coming from the ROI were considered. Next, a sampling interval was chosen (1-100 ms) and positive or negative events were counted within consecutive time bins. The positive or negative event rate (Hz/pixel) at each sampling point was defined as the event count divided by the bin interval *x* total number of pixels constituting the ROI.

#### Processing of FC data

Raw data from the FC consisted of temporal series of consecutive frames. Frame projections for a given recording were generated by calculating the pixel by pixel average grayscale value across all frames. ΔF/F_0_ fluorescence traces for a given ROI (see below) were calculated as follows: first, the average grayscale value within the ROI was calculated at each frame (*F*); next, a baseline value (*F_0_*) was extracted as the 5^th^ percentile of the *F* distribution and subtracted from *F*. Finally, the resulting ΔF was divided by the baseline, detrended with a 2^nd^ order polynomial and multiplied by 100, to obtain the % ΔF/F_0_.

#### Extraction of ROIs and FOV registration

Different methods were used to segment ROIs corresponding to putative neurons. For segmenting GCaMP FC recordings we used an adaptation of a constrained non-negative matrix factorization approach designed for single-photon calcium imaging data (CNMF-E^20^) and implemented in MATLAB^21,22^. For segmenting high resolution event count distributions obtained from GCaMP recordings (for the spontaneous reactivation sequences) we used an online implementation of Cellpose-SAM^23^, based on a foundation model for biological segmentation. Finally, ROIs from BeRST1 fluorescence were manually segmented. Once segmented in one camera, the same ROIs were matched in the other camera via FOV registration. Briefly, EC pixels were mapped onto FC pixels (or vice-versa) by using a two-step process. First, a coarse alignment was performed by applying a rotation followed by cropping and resampling to approximately match the two FOVs at the same pixel resolution. Next, fine alignment was achieved using feature-based image registration using the Scale-Invariant Feature Transform (SIFT). Images were normalized and smoothed with a Gaussian filter, and SIFT key points and descriptors were extracted from both images using identical parameters. Descriptor matching was performed using nearest-neighbor matching, and matched key points were further filtered based on spatial proximity. An affine transformation mapping one image onto the other was then estimated using the MATLAB function *estimateGeometricTransform2D*. The outcome of the registration procedure was visually inspected for every FOV, by looking at overlaid images. Following registration, ROI pixels could be mapped between EC and FC images, allowing a one-to-one correspondence to be established between ΔF/F_0_ fluorescence traces and event rates from the same putative neurons.

#### Detection and quantification of GCaMP fluorescence transients

ΔF/F_0_ fluorescence traces for automatically segmented ROIs from GCaMP spontaneous recordings were processed for fluorescence transients detection as follows. Peaks exceeding 10% ΔF/F_0_ with a peak width > 0.5 s and separated by more than 1s were considered as candidate fluorescence transients. A time window of [-1,+2] s around the peak fluorescence was considered for analysis of the event rate - from the corresponding ROI in the EC FOV - and of the ΔF/F_0_ time derivative (the difference of ΔF/F_0_ between consecutive frames, divided by the frame interval). Peak event rate and peak ΔF/F_0_ time derivative were considered as the maxima (or the minima, in case of negative events) within the [-1, +2] s interval around the peak fluorescence.

#### Data rate calculation

Since the number of events depends on activity levels, the average data rate per neuron from the EC was calculated from empirical data. Briefly, for each segmented neuron from GCaMP recordings, an event rate was calculated by considering the total number of events detected per ROI divided by the recording length. Next, for each ROI a data rate was calculated by multiplying the event rate by 8 bytes, as each event is encoded by 8 bytes. For comparison with the FC, we assumed that the same ROI was imaged at increasing frame rates, and estimated the data rate as ‘*total number of pixels within the ROI*’ x ‘*frame rate*’ x ‘*2 bytes*’, as brightness levels at each pixel in the FC are encoded by 2 bytes (16 bits).

#### Detection of spontaneous activation sequences in vitro

Spontaneous activation sequences from GCaMP fluorescence were detected from EC data as follows. First, in order to identify putative active neurons within a recording, event distributions across pixels were used as input for automatic ROI segmentation, using the Cellpose-SAM^23^ algorithm. Next, for each ROI an event rate was quantified as described above, using 1 ms time bins. A population rate was calculated as the smoothed (0.5 s moving mean) average of the positive event rate for each cell, after normalizing it between 0 and 1. Peaks in the population rate (termed *population bursts*) were identified as those exceeding 1 standard deviation of the trace, with a minimum peak distance of 2 s and a minimum duration of 0.5 s. For each population burst, the activation time of each ROI relative to the burst center was calculated within an interval of ±1.5 *x* burst width, as the time of the peak event rate of the ROI within that interval. Peaks smaller than 2 times the standard deviation of the trace within the interval were not considered for determining the ROI activation order. A median activation order for each ROI across all population bursts was calculated by sorting the median activation times (excluding the invalid ones) and used for sorting ROIs for visualization. Statistical significance of the activation order for each recording was assessed by quantifying the Spearman correlation between the median activation times on random halves of the trials (*C_emp_*) and comparing this empirical value with 100 random shuffles were the correlation was calculated after randomly shuffling the ROI identities (*C_rand_*) in one of the random halves. A *p* value was calculated as the fraction of *C_rand_* higher than *C_emp_*.

### Statistics

No formal methods were applied to predetermine sample sizes and adopted sample sizes were similar to those used by similar studies. No randomization of experimental sessions was performed and no blinding to experimental conditions was implemented during the analysis. All statistical comparisons were performed using nonparametric tests, unless otherwise stated. The tests were two-tailed, unless otherwise stated.

## Data and Code Availability

Data and code used in this study are available from the corresponding author upon reasonable request.

## Acknowledgements

We thank members of the Yartsev laboratory for fruitful discussions and help with the project. In particular, we thank Hesper Chen for help during preliminary experiments and Omkar Ghenand for support in exploring EC biases and FOV registration algorithms. We additionally thank Marisol Navarro for experimental guidance during preliminary experiments in the Miller laboratory. We are grateful to the Adesnik Laboratory (UC Berkeley) for their generous sharing of optical components (0.95 NA, 20x Objective).

## Author Contributions

AF conceived the project. AF, KK and MY designed the research. AF and KK build the microscope. DL, AF and KK performed GCaMP experiments. KK performed BeRST1 experiments. AF performed most of the data analyses. Cultures were prepared by KH, MHM, KK and DL with resources from CL, EM and MMY. BeRST1 was provided by KE and EM. The manuscript was written by AF, KK and MMY with input from all the authors.

## References

1. Kim, T. H. & Schnitzer, M. J. Fluorescence imaging of large-scale neural ensemble dynamics. Cell 185, 9–41 (2022).

2. Barth, A. L. & Poulet, J. F. A. Experimental evidence for sparse firing in the neocortex. Trends Neurosci 35, 345–355 (2012).

3. Marblestone, A. H. et al. Physical principles for scalable neural recording. Front Comput Neurosci 7, 137 (2013).

4. Bai, L. et al. Volumetric voltage imaging of neuronal populations in the mouse brain by confocal light-field microscopy. Nat Methods 21, 2160–2170 (2024).

5. Xiao, S. et al. Large-scale deep tissue voltage imaging with targeted-illumination confocal microscopy. Nat Methods 21, 1094–1102 (2024).

6. Wang, Z. et al. Kilohertz volumetric imaging of in vivo dynamics using squeezed light field microscopy. Nat Methods 22, 2194–2204 (2025).

7. Aharoni, D. & Hoogland, T. M. Circuit investigations with open-source miniaturized microscopes: Past, present and future. Front Cell Neurosci 13, 1–12 (2019).

8. Gallego, G. et al. Event-Based Vision: A Survey. IEEE Trans Pattern Anal Mach Intell 44, 154–180 (2022).

9. Cao, R., Galor, D., Kohli, A., Yates, J. L. & Waller, L. Noise2Image: noise-enabled static scene recovery for event cameras. Optica 12, 46 (2025).

10. Cabriel, C., Monfort, T., Specht, C. G. & Izeddin, I. Event-based vision sensor for fast and dense single-molecule localization microscopy. Nat Photonics 17, 1105–1113 (2023).

11. Howell, J., Hammarton, T. C., Altmann, Y. & Jimenez, M. High-speed particle detection and tracking in microfluidic devices using event-based sensing. Lab Chip 20, 3024–3035 (2020).

12. Moeys, D. P. et al. A Sensitive Dynamic and Active Pixel Vision Sensor for Color or Neural Imaging Applications. IEEE Trans Biomed Circuits Syst 12, 123–136 (2018).

13. Guo, R. et al. EventLFM: event camera integrated Fourier light field microscopy for ultrafast 3D imaging. Light Sci Appl 13, (2024).

14. Chen, T.-W. et al. Ultrasensitive fluorescent proteins for imaging neuronal activity. Nature 499, 295–300 (2013).

15. Lee, A. K. & Wilson, M. A. Memory of sequential experience in the hippocampus during slow wave sleep. Neuron 36, 1183–1194 (2002).

16. Foster, D. J. & Wilson, M. A. Reverse replay of behavioural sequences in hippocampal place cells during the awake state. Nature 440, 680–683 (2006).

17. van der Molen, T. et al. Preconfigured neuronal firing sequences in human brain organoids. Nat Neurosci 29, 123–135 (2026).

18. Ikegaya, Y. et al. Synfire Chains and Cortical Songs: Temporal Modules of Cortical Activity. Science (1979) 304, 559–564 (2004).

19. Huang, Y.-L., Walker, A. S. & Miller, E. W. A Photostable Silicon Rhodamine Platform for Optical Voltage Sensing. J Am Chem Soc 137, 10767–10776 (2015).

20. Zhou, P. et al. Efficient and accurate extraction of in vivo calcium signals from microendoscopic video data. Elife 7, e28728 (2018).

21. Liberti, W. A., Schmid, T. A., Forli, A., Snyder, M. & Yartsev, M. M. A stable hippocampal code in freely flying bats. Nature 604, 98–103 (2022).

22. Forli, A. & Yartsev, M. M. Hippocampal representation during collective spatial behaviour in bats. Nature 621, 796–803 (2023).

23. Pachitariu, M., Rariden, M. & Stringer, C. Cellpose-SAM: superhuman generalization for cellular segmentation. bioRxiv 2025.04.28.651001 (2025).

